# Cannabis Vapour Exposure Alters Neural Circuit Oscillatory Activity In A Neurodevelopmental Model Of Schizophrenia: Exploring The Differential Impact Of Cannabis Constituents

**DOI:** 10.1101/2021.04.28.441799

**Authors:** Bryan W. Jenkins, Shoshana Buckhalter, Melissa L. Perreault, Jibran Y. Khokhar

**Affiliations:** Department of Biomedical Sciences, University of Guelph, Guelph, Ontario, Canada; Department of Molecular and Cellular Biology, University of Guelph, Guelph, Ontario, Canada

**Keywords:** cannabinoid, oscillations, preclinical, psychosis, electomes, oscillopathies

## Abstract

Cannabis use is highly prevalent in patients with schizophrenia and worsens the course of the disorder. To understand the causal impacts of cannabis on schizophrenia-related oscillatory disruptions, we herein investigated the impact of exposure to cannabis vapour (containing delta-9-tetrahydrocannabinol [THC] or balanced THC and cannabidiol [CBD]) on oscillatory activity in the neonatal ventral hippocampal lesion (NVHL) rat model of schizophrenia. Male Sprague Dawley rats underwent NVHL or sham surgeries on postnatal day 7. In adulthood, electrodes were implanted targeting the cingulate cortex (Cg), the prefrontal cortex (PFC), the dorsal hippocampus (HIP), and the nucleus accumbens (NAc). Local field potential recordings were obtained following exposure to two strains of vapourized cannabis flower (with ~10% THC or ~10% balanced THC:CBD) in a cross-over design with a two-week wash-out period between exposures. Compared to controls, NVHL rats had reduced baseline gamma power in the Cg, dHIP, and NAc, and reduced high-gamma coherence between the dHIP-Cg. THC-only vapour broadly suppressed oscillatory power and coherence, even beyond the baseline suppressions observed in NHVL rats. Balanced THC:CBD vapour appeared to ameliorate the THC-induced impacts on power and coherence in both sham and NVHL rats. For NVHL rats, THC-only vapour also normalized the baseline dHIP-Cg high-gamma coherence deficits. NHVL rats also demonstrated a 20ms delay in dHIP theta to high-gamma phase coupling, which was ameliorated by both exposures in the PFC and NAc. In conclusion, THC-only cannabis vapour suppressed oscillatory activity in NVHL and sham rats, while balanced THC:CBD vapour may ameliorate some of these effects.

## Introduction

The lifetime rate of cannabis use by patients with schizophrenia is reportedly as high as 80%, with as many as 50% of patients dually diagnosed with cannabis-use disorder and schizophrenia ^1^. Cannabis use is associated with a worsened prognosis through treatment non-compliance, increased rates of hospitalization and relapse, and by exacerbating symptoms ^1–3^. Considering the variety of effects produced by cannabis constituents ^4^, cannabis use may produce varying outcomes for patients depending on the constituent profile of the cannabis used and the chosen administration route ^5–7^. The varying effects of cannabis arise from its phytochemical diversity; with over 113 constituents, or ‘cannabinoids’, their relative composition dictates the effects produced ^4^. Due to their abundance and relevance for biomedical research, scientific investigation has mainly focused on two cannabinoids: Δ9-tetrahydrocannabinol (THC) and cannabidiol (CBD). THC is the main psychoactive constituent that binds with endogenous cannabinoid type 1 receptors (CB1Rs) to produce the rewarding and psychotomimetic effects ^4, 8–10^. In contrast, CBD is non-psychoactive and acts as a negative allosteric modulator of CB1Rs, while also being a partial agonist of both cannabinoid type 2 and dopamine type 2 receptors, and an agonist of serotonin 1A receptors ^11–15^. Moreover, CBD is suggested to have antipsychotic potential and oppose the psychoactive effects of THC ^16–19^. Additionally, the independent effects of these constituents differ from the effects produced when co-administered, with these effects also being affected by administration route ^20–22^.

THC modulates brain signals and behaviour in healthy individuals, as well as in individuals with schizophrenia. When administered intravenously (IV), THC produces transient states of psychosis, supporting the psychotomimetic properties of THC ^23–25^. IV THC-induced psychosis also correlates with reduced oscillatory coherence, specifically in the gamma frequency band ^26^. Furthermore, IV THC administered in patients with schizophrenia reduces spectral power below 27Hz, and enhances power above 27Hz ^27^, while also transiently exacerbating symptoms in patients. Specifically, IV THC transiently increases the positive, negative, and cognitive symptoms of schizophrenia, including learning, memory and perceptual deficits ^7, 28^. This is contrasted by evidence that synthetic THC (dronabinol) reduces psychotic symptoms in patients ^29, 30^. The highly variable composition of the cannabis strains, THC concentrations and the routes of administration may explain the differential effect on symptoms and correlated neural activity.

Recently Wall et al. ^5^ used magnetic resonance imaging (MRI) to measure functional connectivity changes in the default mode network (DMN, measured as positive connectivity in the posterior cingulate cortex), executive control network (ECN, negatively correlated with DMN activity and measured as negative connectivity in the posterior cingulate cortex), and salience network (SAL, measured as positive connectivity within the anterior insula), after healthy controls were administered either vapourized placebo cannabis, cannabis with THC and CBD (8mg and 10mg), or cannabis without CBD (THC: 8mg). Cannabis vapour with, or without, CBD reduced DMN connectivity, relative to placebo, while not significantly affecting ECN connectivity. Only vapour without CBD impacted SAL connectivity, reducing it relative to vapour with CBD. Reduced DMN activity correlated with the self-reported psychoactive effects of THC while the vapour with CBD exposure had an ameliorative effect ^5^. Similarly, we previously assessed DMN functional connectivity in patients with co-occurring schizophrenia and cannabis use disorder, as well as otherwise healthy controls, using MRI scans taken before and after subjects smoked cannabis or ingested THC (15mg) ^31^. Before cannabis exposure, patients exhibited DMN hyperconnectivity compared to controls, and this hyperconnectivity was correlated with positive symptom severity; patients also exhibited reduced anti-correlation between DMN and executive control network. Cannabis exposure reduced hyperconnectivity and increased anti-correlation. Working memory performance was also assessed and found to be positively correlated with the magnitude of anti-correlation in controls and patients that were administered cannabis ^31^.

Neural circuit dysfunction is apparent in rodent models of schizophrenia and controls, with and without cannabinoid administration ^32^). Local field potential (LFP) recordings from the neonatal ventral hippocampal lesion (NVHL) rat model of schizophrenia demonstrate reduced theta and beta coherence in the dHIP, with intact coherence in the medial prefrontal cortex (PFC), compared to controls ^33^. Phase-locking of auditory evoked potentials in the temporal cortex of control rats increases with increasing stimulus frequencies, and yet NVHL rats do not exhibit this relative increase; thus, NVHL rats exhibit a breakdown of phase-locking in their baseline temporal cortical oscillatory activity ^34^. Enhanced power in the upper theta and beta frequency range, as well as reduced delta and high-gamma power, is also apparent in Wistar rats selectively bred to exhibit maximal disruptions after undergoing post-weaning social isolation and sub-chronic ketamine administration, used to model schizophrenia ^35^.

In experimentally-naïve rats, LFP recordings from the hippocampus and entorhinal cortex demonstrate that intraperitoneal administration of CB1R agonist CP 55,940 reduces evoked theta and gamma power; this was reproduced in humans administered IV THC and whole-brain electroencephalography (EEG) ^36^. We previously published data demonstrating acute THC vapour exposure in rats suppresses LFP gamma power and coherence in the dorsal striatum, the PFC, and the orbitofrontal cortex ^37^. Considering the opposing effects of THC and CBD on neural circuit activity in patients and models of schizophrenia and in healthy controls, we herein aimed to 1) detect baseline differences in LFPs from the PFC, cingulate cortex (Cg), dHIP, and nucleus accumbens (NAc) of NVHL rats and sham-surgery (sham) controls; and 2) to assess changes in LFPs in NVHL and sham rats resulting from exposure to THC-containing cannabis vapour with and without CBD. We hypothesize that the presence of CBD in cannabis vapour will ameliorate THC-induced reductions in corticolimbic oscillatory activity and may rescue baseline deficits in the NVHL rat.

## Methods

### Animals

Ten male Sprague Dawley rat pups with lactating dam were purchased from Charles River. Rats were housed in a colony room maintained on a 12-hour light:dark cycle with *ad libitum* access to food and water. Prior to the start of experiments, rats were habituated to the experimental environment for two minutes daily and for a total of five days. All treatments were performed during the light phase of the 12-hour reverse light:dark cycle. All procedures complied with the guidelines described in the Guide to the Care and Use of Experimental Animals (Canadian Council on Animal Care, 1993) and set by the Animal Care Committee at the University of Guelph.

### The neonatal ventral hippocampal lesion rat model of schizophrenia

The NVHL rat was produced as previously described ^38, 39^; the NVHL rat is considered a valuable heuristic model of schizophrenia and co-occurring substance use ^40^, including in the context of cannabis use and schizophrenia ^41, 42^. This neurodevelopmental model involves bilaterally lesioning the ventral hippocampi of rat pups using ibotenic acid. On postnatal day 7 (PND7) rat pups were anaesthetized via hypothermia, injected with either 0.3μL of ibotenic acid (for the NVHL group) or artificial cerebrospinal fluid (aCSF, for the sham group) into the ventral hippocampus (anterior-posterior [AP]: −3 mm; medial-lateral [ML]: ± 3.5 mm; dorsal-ventral [DV]: −5mm) at a rate of 0.1μL/min over three minutes followed by a two-minute wait period. Animals were housed with mom in polyethylene cages until weaning, and pair-housed until the electrode implantation surgery.

### Electrode implantation surgeries

Electrode implantation was performed as previously described ^43^ (Figure 1A). Custom-built, 10-channel electrode microarrays were constructed using Delrin templates and polyimide-insulated stainless-steel wires (A-M Systems, 791600, 0.008”) attached to a single-row connector. All arrays used had an electrode impedance of less than 2MΩ. Rats were anesthetized with isoflurane (5%), and electrode arrays were implanted with bilateral electrodes into the medial PFC (AP: +3.24 mm, ML: ±0.6 mm, DV: −3.8 mm), the Cg (AP: +1.9 mm, ML: ±0.5 mm, DV: −2.8 mm), the CA1 region of the dHIP (AP: −3.5mm, ML: ±2.5mm, DV: −2.6mm), and the NAc (AP: +1.9 mm, ML: ±1.2 mm, DV: −6.6 mm). Animals recovered in their home cage for a minimum of 7 days prior to experimentation.

**Figure 1.**
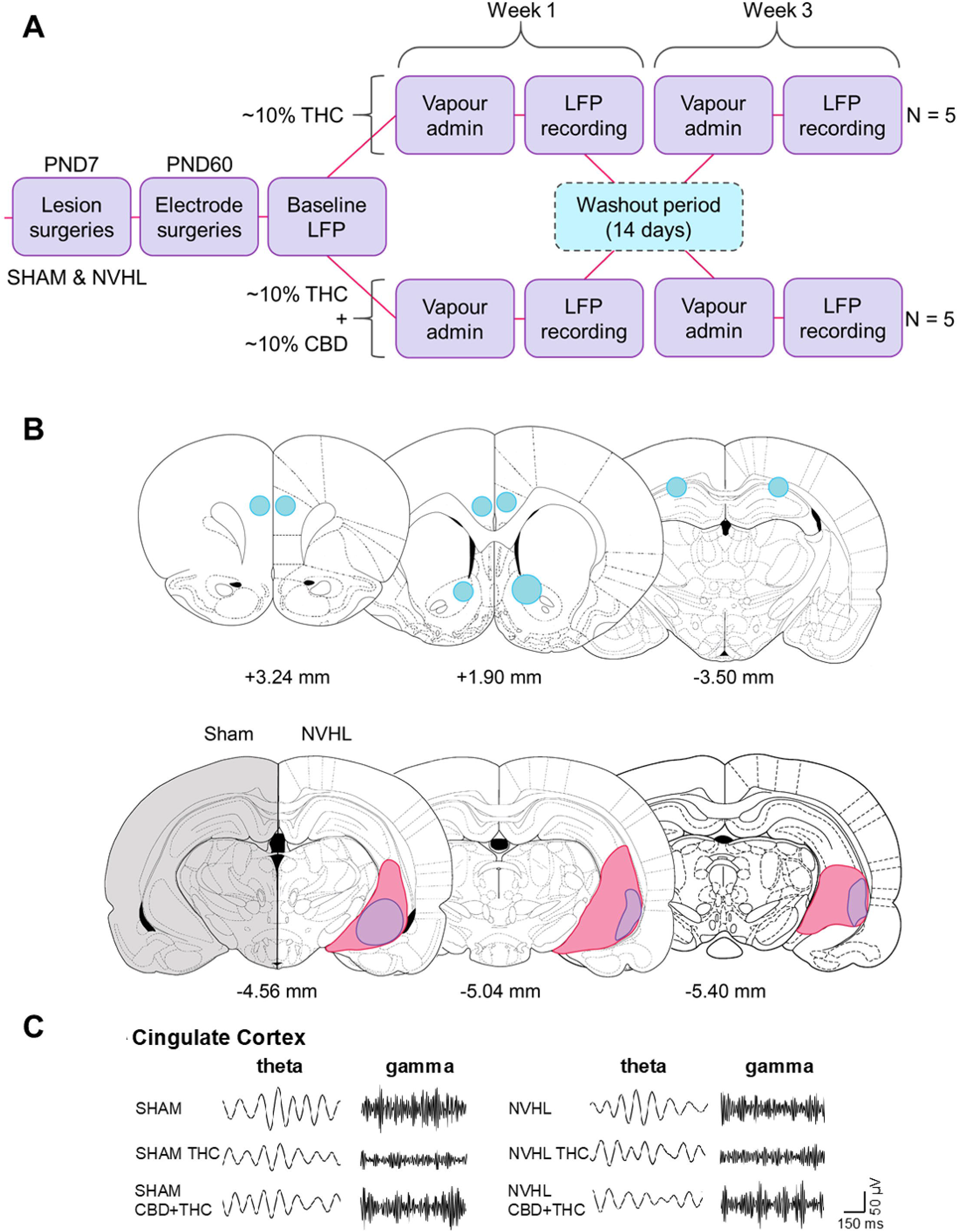
Experimental design, lesion and electrode verification, and representative local field potential tracing. **A)** Cross-over design with a two-week wash-out period between exposures **B)** Electrode placements (top) and lesion (bottom) showing area encompassing electrode termini (blue) in the PFC (left), Cg + NAc (middle), and HIP (right), as well as lesion extent in all NVHL rats (purple represents size of smallest and pink represents size of largest lesions), compared to sham controls. **C)** Representative tracings from the Cg showing changes in theta and gamma frequencies at baseline and after cannabis vapour exposure for both sham and NVHL rats.

### Vapourized THC/CBD administration

Cannabis flower vapour administration was performed using a customized OpenVape apparatus with a Utillian 420 dried herb vapourizer ^44–46^. A crossover design was used as previously performed in human subjects and in our study with rats ^20, 37^. Rats were randomly assigned to two groups (N = 5/group) for exposure to either cannabis vapour with 8-18% THC and 0% CBD (Aurora Summer Fling; THC Vapour) or 4-11% THC and 8.5-15.5% CBD (Solei Balance: THC:CBD Vapour). Ground cannabis flower was vapourized at 226 °C and channelled into an empty mouse cage (40 × 20 × 20 cm).

### Electrophysiology

LFP recordings (Wireless 2100-system, Multichannel Systems) were performed in awake, freely moving rats exploring clear plexiglass boxes (45 cm × 45 cm × 45 cm). Baseline recordings were collected 24 hours before experimentation and immediately before cannabis vapour exposure, for 30 min at a rate of 1000 samples/s. MATLAB (MathWorks) routines from the Chronux software package were used to analyze the spectral power of each brain region, as well as coherence and theta-gamma cross-correlation between brain regions. Five-minute epochs were used, with epochs segmented, detrended, denoised, and low-pass filtered to remove frequencies greater than 100 Hz. Continuous multi-taper spectral power for data normalized to total spectral power and coherence (tapers=[5 9]) was calculated for delta (1-4 Hz), theta (>4-12 Hz), beta (>12-30 Hz), low gamma (>30-60 Hz), and high gamma (>60-100 Hz).

### Histology

After experimentation, rats were euthanized by carbon dioxide + isoflurane before their brains were extracted and flash frozen. Brains were subsequently sectioned (at 40 mm) using a cryostat, mounted on slides, and stained with thionin. The Cg, PFC, dHIP, and NAc were examined microscopically to confirm electrode placement and bilateral lesioning. Rats with unilateral or extrahippocampal lesions or misplace electrodes would be removed from the analyses.

### Data analysis

Analysis was performed on one-minute time bins with data curves presented as normalized data with jack-knife estimates of standard error of the mean (SEM). Data was log-transformed to better exhibit group differences. Prior to all analyses, normality was assessed using the Shapiro-Wilk test. Sham or NVHL group comparisons were performed using a repeated measures ANOVA with treatment as the within-subjects factor, followed by paired t-tests. A Student’s t-test was used to compare NVHL exposure groups to baseline sham data. Computations were performed using IBM SPSS 24 software and are expressed as means ± SEM or percent change from sham baseline ± SEM.

## Results

Lesions and electrode placements were verified microscopically in NVHL and sham rats. All NVHL rats showed bilateral lesions of the ventral hippocampus (Figure 1B indicates the minimum and maximum lesion sizes observed), and the electrodes were verified to be in the correct locations (Figure 1B). Representative theta and gamma oscillations obtained from the Cg are shown in Figure 1C.

Baseline gamma deficits in spectral power exist in NVHL rats across all brain regions, when compared to sham rats, and these deficits are selectively modified by cannabis vapour exposure (Figure 2). In the Cg, the only baseline deficit apparent between NVHL and sham rats was with high-gamma power (p=0.021, Figure 2B). THC:CBD-vapour exposure attenuated this high-gamma baseline deficit in NVHL rats (Figure 2B). THC-only vapour exposure reduced Cg beta (p=0.001) and low-gamma power (p=0.019) only in NVHL rats, an effect that was also ameliorated after THC:CBD-vapour exposure. Both exposures enhanced delta, and reduced theta, power in the Cg of all rats [sham delta: F(2,14)=6.7, p=0.009, theta: F(2,14)=9.2, p=0.003; NVHL delta: F(2,14)=3.5, p=0.059, theta: F(2,14)=7.5, p=0.008] (Figure 2B). In the PFC, baseline power was similar between NVHL and sham rats across all frequency bands (Figure 2C). For NVHL rats, THC-only vapour exposure suppressed PFC beta power while the THC:CBD-vapour exposure ameliorated this effect [NVHL beta: F(2,14)=8.9, p=0.003]. For all rats, both exposures reduced theta power, and delta power to a lesser degree [sham delta: F(2,14)=5.6, p=0.016, theta: F(2,14)=32.3, p<0.001; NVHL delta: F(2,14)=3.3, p=0.067, theta: F(2,14)=20.6, p<0.001], while THC-only vapour exposure reduced both low- and high-gamma power; THC:CBD-vapour exposure did not have any effect [sham low gamma: F(2,14)=6.7, p=0.009, high gamma: F(2,14)=4.4, p=0.032; NVHL low gamma: F(2,14)=4.0, p=0.042].

**Figure 2.**
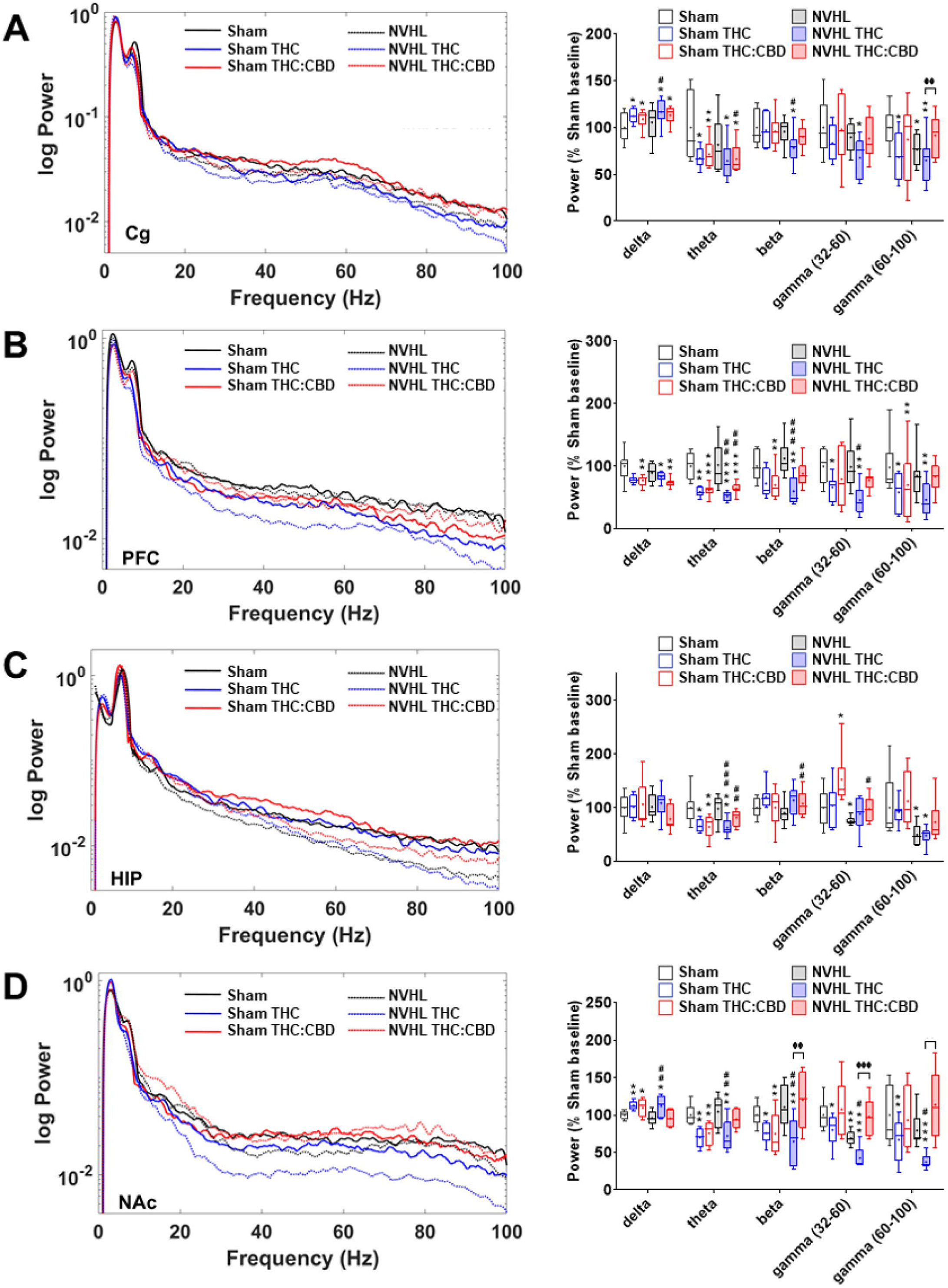
NVHL rats exhibit reduced baseline spectral power compared to sham rats, and cannabis vapour differentially modulates this dysfunction. **A)** Spectral power tracing (left) and quantification (right) in the Cg showing baseline deficits in high-gamma power for NVHL rats compared to sham rats; THC-only vapour reduced Cg beta and low-gamma power only in NVHL rats, an effect that was ameliorated after THC:CBD-vapour exposure; THC:CBD-vapour exposure also attenuated the high-gamma baseline deficit in NVHL rats; both exposures enhanced delta, and reduced theta, power in the Cg of all rats; **B)** In the PFC,THC-only vapour exposure suppressed PFC beta power while the THC:CBD-vapour exposure ameliorated this effect in NVHL rats; for all rats, both exposures reduced theta power, and delta power to a lesser degree, while THC-only vapour exposure reduced both low- and high-gamma power; THC:CBD-vapour exposure did not have any effect; **C)** In the dHIP, NVHL rats showed baseline deficits in low- and high-gamma power, compared to sham rats whereas THC:CBD-vapour exposure appeared to marginally improve baseline deficits; both exposures did not affect delta power and robustly suppressed theta power in NHVL and sham rats. **D)** In the NAc, NVHL rats showed baseline deficits in low-gamma power, compared to sham controls; THC-only vapour exposure reduced spectral power across all other frequencies in both NVHL and sham rats; THC:CBD exposure ameliorated these effects across all frequencies for NVHL rats, and selectively within the gamma frequency band for sham rats. □P<0.05; □□P<0.01 compared to sham control baseline; ##P<0.05 compared to NVHL rat baseline; φP<0.05 compared THC-only treated rats to THC:CBD treated rats.

In the dHIP of NVHL rats, baseline deficits in low- (p=0.044) and high-gamma (p=0.047) power were observed, compared to sham controls (Figure 3A); these deficits appeared to be marginally improved after the THC:CBD-vapour exposure, while the THC-only vapour exposure did not show any differences from baseline. Similarly, both exposures did not affect delta power in NHVL or sham rats. Both exposures, however, robustly suppressed theta power in all rats [sham theta: F(2,14)=32.3, p<0.001; NVHL theta: F(2,14)=20.6, p<0.001]. In the NAc, NVHL rats had baseline deficits in low-gamma power, compared to sham controls (p=0.001, Figure 3B). THC-only vapour exposure increased NAc delta power for all rats while THC:CBD exposure enhanced delta power for sham rats only [sham delta: F(2,14)=4.7, p=0.027; NVHL delta: F(2,14)=6.6, p=0.010]. Notably, THC-only vapour exposure reduced spectral power across all other frequencies in both NVHL and sham rats, while THC:CBD exposure ameliorated these effects across all frequencies for NVHL rats, and selectively within the gamma frequency band for sham rats [sham theta: F(2,14)=13.2, p=0.001; low gamma: F(2,14)=4.8, p=0.026; high gamma: F(2,14)=9.9, p=0.002; NVHL theta: F(2,14)=7.8, p=0.005; beta: F(2,14)=10.3, p=0.002; low gamma: F(2,14)=8.0, p=0.005; high gamma: F(2,14)=15.6, p=0.001].

**Figure 3.**
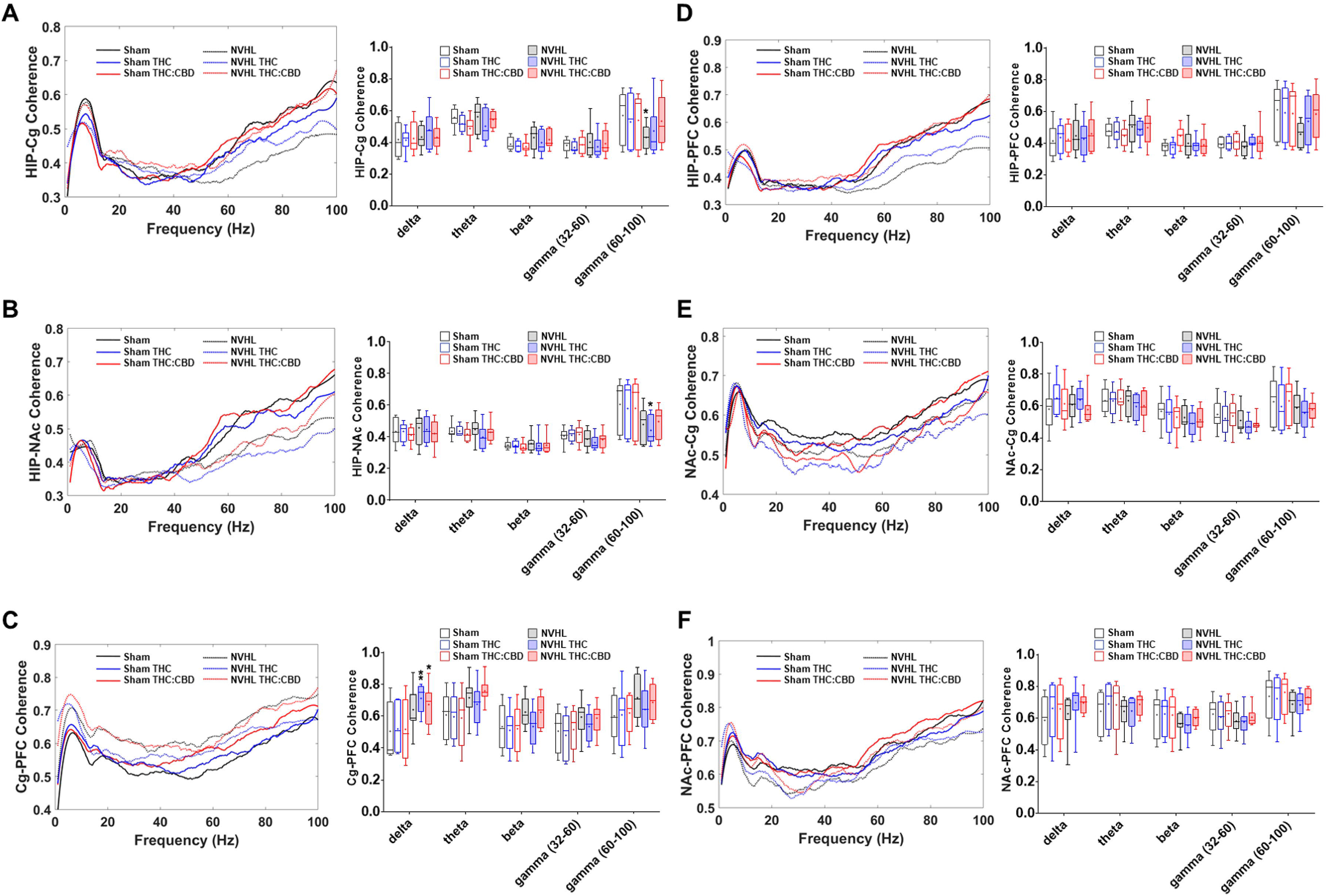
NVHL rats exhibit reduced baseline coherence within and between cortical and limbic regions, with variable impacts of cannabis vapour. **A)** Both vapour exposures enhanced dHIP-Cg high-gamma coherence in the NVHL rats to levels that were no longer different from sham baseline coherence; **B)**, NVHL rats exhibited a trend toward reduced dHIP-NAc high-gamma coherence, compared to controls, and THC-only exposure exacerbated this deficit; **C)** Both types of cannabis-vapour exposures enhanced Cg-PFC delta coherence in NVHL rats to levels above baseline coherence in sham rats; Cannabis vapour exposure did not alter **D)** dHIP-PFC, **E)** NAc-Cg, or **F)** NAc-PFC coherence. □P<0.05; □□P<0.01 compared to sham control baseline.

Measures of oscillatory coherence between the target regions revealed that baseline differences exist between NVHL and sham rats and are modifiable by cannabinoid exposure (Figure 4). Cannabis vapour exposure did not alter dHIP-PFC (Figure 4B), NAc-Cg (Figure 4D), or NAc-PFC (Figure 4E) coherence. Both types of cannabis-vapour exposures enhanced Cg-PFC delta coherence in NVHL rats to levels above baseline coherence in sham rats (THC: p=0.005; THC:CBD: p=0.039; Figure 4F). Between limbic regions dHIP-NAc, a trend toward the NVHL rats exhibiting reduced high-gamma coherence was observed, when compared to controls (p=0.060, Figure 4C). THC-only exposure exacerbated the high-gamma coherence deficit between dHIP-NAc (p=0.018). NVHL rats exhibited reduced high-gamma power between the dHIP-Cg, compared to controls (p=0.041, Figure 4A). Both vapour exposures enhanced HIP-Cg high-gamma coherence in the NVHL rats to levels that were no longer different from sham baseline coherence.

**Figure 4.**
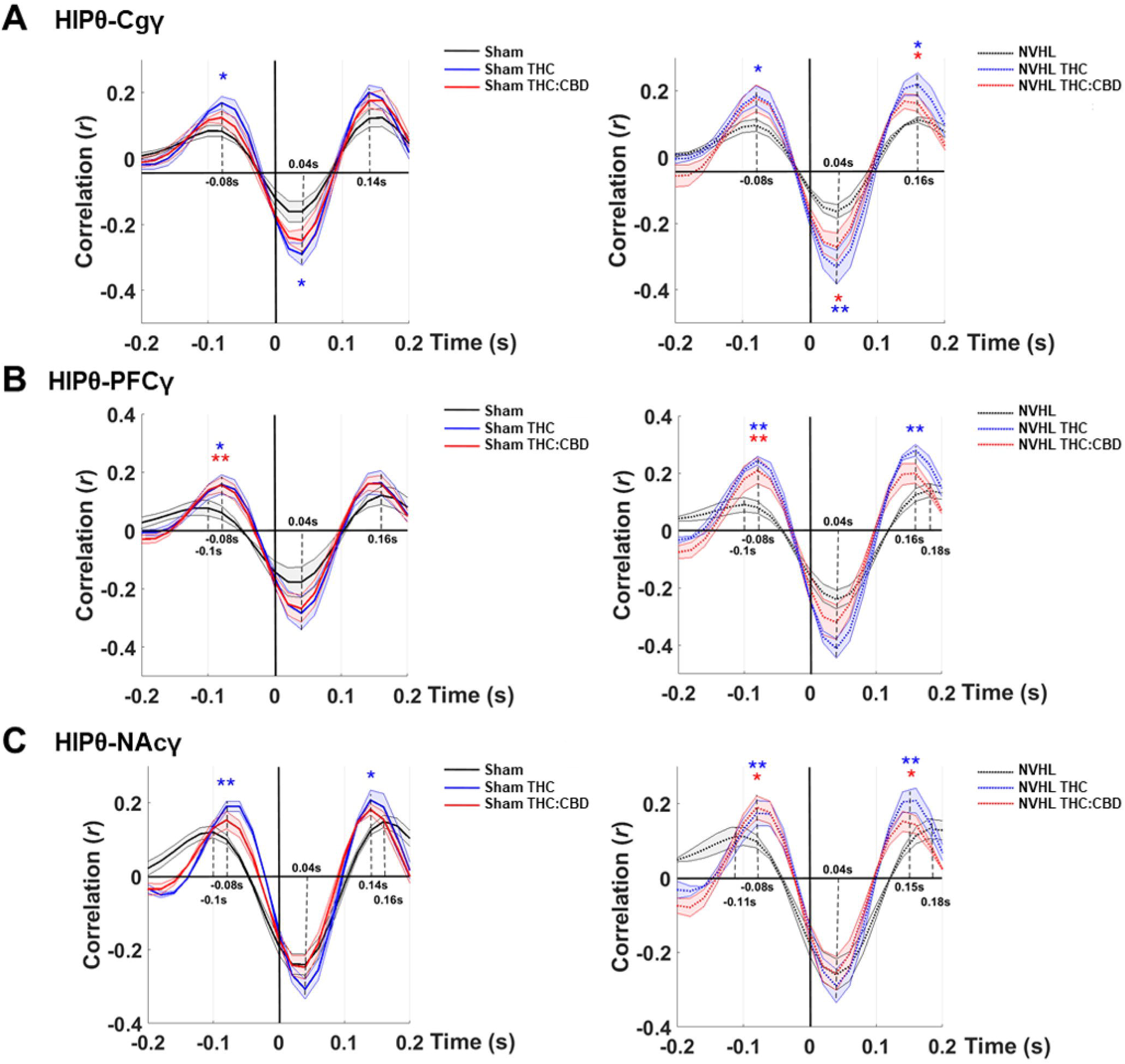
NVHL rats exhibit baseline deficits in phase coupling between dHIP theta and Cg, PFC, or NAc gamma, and cannabis vapour exposure selectively ameliorates this delay. **A-C)** NVHL rats exhibited a 20ms delay in coupling of dHIP theta to high-gamma in the Cg, PFC, and NAc; both vapour exposures ameliorated this delay in the **B)** PFC and **C)** NAc, but not in the **A)** Cg; For all rats, the THC-only vapour exposure also enhanced the dHIP theta-gamma coupling strength in all regions, an effect that was also observed to a lesser extent after THC:CBD-vapour exposure; NVHL rats exhibited baseline theta-gamma coupling strength similar to sham rats, for all pairwise comparisons. □P<0.05; □□P<0.01 compared to sham control baseline.

Cross-correlation analysis revealed that baseline deficits in phase coupling between dHIP theta and Cg, PFC, or NAc gamma in the NVHL rats is sensitive to the effects of cannabis vapour exposure (Figure 4). Comparing theta-gamma coupling strength between NVHL and sham rats revealed similar baseline values for all pairwise comparisons; however, compared to sham rats, NVHL rats exhibited a 20ms delay in coupling of dHIP theta to high-gamma in all other regions. Both vapour exposures ameliorated this delay in the PFC and NAc (Figure 4D, F) but not in the Cg (Figure 5B). For all rats, the THC-only vapour exposure also enhanced the dHIP theta-gamma coupling strength in all regions. This effect was also observed to a lesser extent after THC:CBD-vapour exposure.

**Figure 5.**
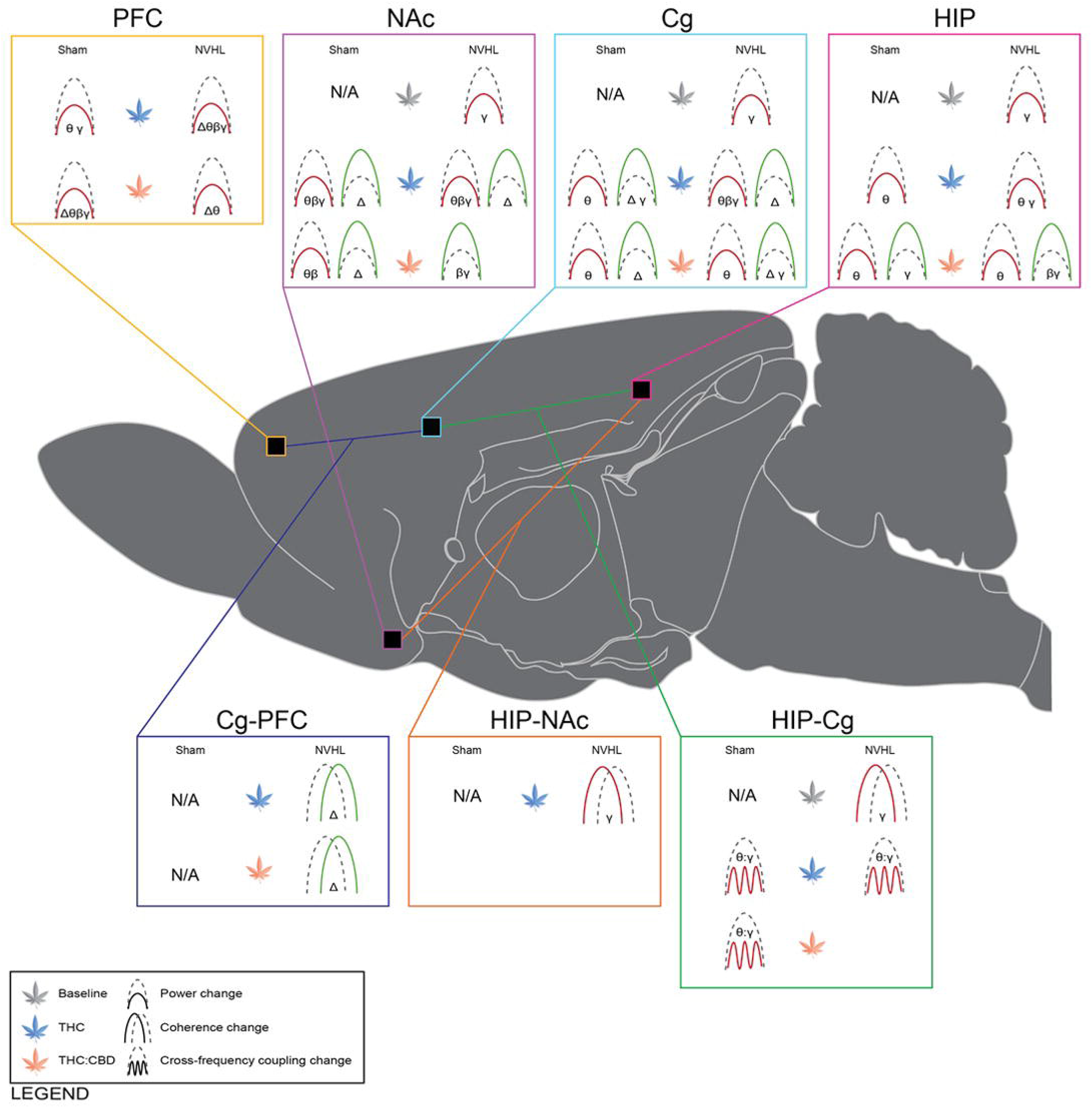
Graphical summary of power, coherence and phase coupling changes associated with NVHL lesions and cannabis vapour exposure. Cannabis vapour exposure produced constituent- and region-dependent disruptions in neural circuit oscillatory activity, often reducing theta, beta, and gamma power while enhancing delta power. Cannabis vapour exposure also enhanced coherence in cortical regions while reducing coherence and phase coupling between limbic regions. Cannabis vapour exposure further exacerbated baseline NAc, Cg, HIP gamma power deficits apparent in NVHL rats, compared to sham controls. PFC = prefrontal cortex; NAc = nucleus accumbens; Cg = cingulate cortex; HIP = hippocampus. Δ = delta frequency band; θ = theta frequency band; β = beta frequency band; γ = gamma frequency band. Green = increase; red = decrease. Icons represent comparisons to sham baseline values and thus ‘N/A’ is used for baseline sham findings.

## Discussion

This study assessed differences in baseline oscillatory activity between NVHL and sham rats, as well as changes in oscillatory activity following exposure to THC-containing cannabis vapour with, or without, similar amounts of CBD. Compared to sham rats, NVHL rats exhibit baseline differences in gamma power and coherence, matching observations in patients with schizophrenia. NVHL rats also demonstrate no difference in baseline delta, theta, and beta power, compared to controls, and enhanced Cg-PFC delta coherence. Whole-brain EEG recordings of oscillatory activity in patients with schizophrenia demonstrate that, compared to controls, patients have reduced evoked theta (3-7Hz), alpha (8-12Hz), beta (12-30Hz), and gamma (30-100Hz) activity associated with deficits in working memory ^47^. Reductions in evoked beta and gamma power are also associated with P50 gating deficits, a hallmark feature in EEG traces from patients with schizophrenia that relates to symptomatic sensory processing deficits ^48^. Patients with schizophrenia also exhibit dysfunctional prefrontal-cortical gamma band activity during high and low cognitive demands that correlate with psychotic symptoms ^49^. Finally, slow-wave delta, theta, alpha, and beta frequencies are enhanced in patients with schizophrenia, compared to controls, along with enhanced delta and alpha coherence between various brain regions ^50–52^. Previous studies using various rodent models of schizophrenia report reduced theta power (in the HIP and prelimbic cortex of DISC1 knock-out mice) and reduced theta and beta coherence (in the dHIP of NVHL rats) ^33, 53^. Variance between our results and existing preclinical literature could be due to the use of different rodents to produce these models (C57BL/6J mice and Long Evans rats, respectively, in the cited papers), while the variance between clinical and preclinical measures may be explained by the inherent differences in oscillatory activity that exist between rodents and humans ^54^.

For all NVHL and sham rats in our study, THC-only vapour exposure had an overall suppressive effect on spectral power across all frequencies except delta, with the THC:CBD-vapour exposure counteracting some of these effects. Interestingly, the restorative effect of THC:CBD-vapour exposure seems most pronounced in NVHL rats, compared to sham controls. After both exposures, Cg-PFC coherence was enhanced, rescuing baseline deficits in the NVHL rats, whereas coherence between limbic regions was exacerbated by THC-only exposure. Moreover, THC-only vapour exposure strengthened Cg-PFC coherence for all rats. As observed previously in humans administered THC ^26^, we demonstrated that exposure to THC-only vapour suppresses gamma power, and further demonstrated that NVHL rats are susceptible to shifts in power induced by exposure to THC-only vapour ^27^; however, EEG recordings from patients with schizophrenia show increased amplitudes for frequencies greater than 27Hz. Our results indicate that exposure to THC-only vapour reduces spectral power in all frequencies but the delta band. This could be due to the different administration routes ^55^, as we chose an administration protocol with more translational relevance ^56^. It could also be the result of the doses administered in this study, as previous research using the PCP rat model of schizophrenia demonstrated through single-unit recordings in the ventral tegmental area (VTA) before and after two doses of intraperitoneal THC that baseline deficits in the number of active neurons in the VTA of PCP rats are reversed only by exposure to a low dose of THC (0.1mg/kg), while exposure to a higher dose (1mg/kg) did not reverse baseline deficits. Both the low and high dose decreased the number of active neurons in control rats ^57^. PCP rats administered the same dose of THC (1mg/kg) also demonstrate dysfunctional activity in the PFC, evidenced by an absence of the THC-induced disinhibition of neuronal firing observed in controls rats ^58^.

Our data also support past observations that CBD opposes some of the actions of THC ^22^. Infusion of THC (100 ng) directly into the ventral HIP (vHIP) of anaesthetized male Sprague Dawley rats enhances neuronal firing and bursting rates, as well as beta and gamma power, in the VTA; rats that were intracranially infused with CBD (100 ng) or co-infused THC + CBD (100 ng each) into the vHIP demonstrate no difference in oscillatory power when compared to vehicle-treated controls ^22^. While a complete understanding of the mechanism of action for the potential antipsychotic effects of CBD remains elusive, they are mediated in part through the normalization of mediotemporal, mediotemporal-striatal, and prefrontal cortical activity, measured using functional magnetic resonance imaging in patients with schizophrenia before, and after, exposure to CBD; this normalization of brain activity corresponded with improvements in self-reported symptom severity ^59^.

Preclinical evidence also supports the involvement of rescued oscillatory dysfunction in mediating antipsychotic action. Aberrant delta and theta power and coherence was detected in LFPs from the PFC of the methylazoxymethanol rat model of schizophrenia, and this was normalized by chronic antipsychotic administration ^60^. Thus, CBD may also exert antipsychotic effects in part by normalizing aberrant neural circuit oscillatory activity. Additionally, the widespread impact of cannabinoid exposure in schizophrenia may be due in part to the involvement of the endocannabinoid system (eCB) in the etiology of the disorder. Patients with schizophrenia have altered eCB functioning, evidenced by elevated levels of anandamide and post-mortem CB1R expression in patients ^61, 62^. Exposure to exogenous cannabinoids may shift the homeostatic norm in patients such that it acutely restores system function in some regions to levels associated with typical functioning, while concurrently exacerbating dysfunctions in other regions.

The baseline 20ms-delay deficit in phase coupling that we observed in NVHL rats was also ameliorated after both exposures in all regions but the Cg. Impaired theta-gamma phase coupling is apparent in patients with schizophrenia, compared to healthy controls, and is associated with working memory impairments ^63^. Theta-gamma phase coupling was improved in patients with schizophrenia that were using cannabis in addition to nicotine, compared to patients only using nicotine ^64^. Similarly, dual diagnosis patients exhibit, through measuring resting-state functional connectivity, baseline hyperconnectivity in the DMN that corresponds with positive symptom severity and reduced negative correlation between DMN and ECN. After oral THC or smoked cannabis administration, patients exhibit reduced DMN hyperconnectivity and restored negative correlation with the ECN, while the magnitude of the negative correlation further correlated with working memory performance ^31^.

The results of this study reveal that cannabis can have a varying impact on oscillatory activity, and the routes of administration, dose, and compositions of cannabis or synthetic cannabinoids may be partly responsible for the varying cannabis-related outcomes reported in patients with schizophrenia. Importantly, our results posit a question of whether advising patients to consume cannabis with balanced THC:CBD levels could be a viable harm reduction strategy. This study is limited in that the other constituents in the selected cannabis strains were not matched and could also have unknown effects ^65^. Additionally, our selection of a within-subject design means that rats were exposed to cannabis vapour twice and there could be residual effects of prior exposure ^37^. However, for this study we selected a two-week washout period, and we did not observe any significant differences between the baseline recordings and recordings captured before the vapour exposure two weeks later, thus confirming that the washout period duration was sufficient. Lastly, only male rats were used in this study. Although cannabis use in schizophrenia is primarily observed in male patients ^66^, previous studies have suggested that considerable sex differences exist in the oscillatory biomarkers associated with psychopathology ^43^, warranting the assessment of sex as a biological variable in the measures assessed here.

Future investigations will be performed using the same protocol to detect any changes in schizophrenia-like behaviours after acute and chronic cannabis vapour exposure using the NVHL rat, as past investigations indicate that THC-induced reductions in oscillatory activity relate to deficits in sensorimotor gating and working memory ^36 67^; we will also aim to identify which electrophysiological features are related to behavioural deficits ^68^. Additionally, understanding how cessation of cannabis use impacts aberrant oscillatory activity may help further characterize the causal contribution of oscillatory activity to effects of cannabis vapour exposure on the symptoms of schizophrenia.

## Acknowledgements

The authors graciously acknowledge the contributions of M. Asfandyaar Talhat, Bryana Hallam, and Chuyun (Judy) Chen for their assistance with the execution of experiments.

